# The SARS-CoV-2 spike (S) and the orthoreovirus p15 cause neuronal and glial fusion

**DOI:** 10.1101/2021.09.01.458544

**Authors:** Ramon Martinez-Marmol, Rosina Giordano-Santini, Eva Kaulich, Ann-Na Cho, Md Asrafuzzaman Riyadh, Emilija Robinson, Giuseppe Balistreri, Frédéric A. Meunier, Yazi D. Ke, Lars M. Ittner, Massimo A. Hilliard

## Abstract

Numerous enveloped viruses use specialized surface molecules called fusogens to enter host cells^1^. During virus replication, these fusogens decorate the host cells membrane enabling them the ability to fuse with neighboring cells, forming syncytia that the viruses use to propagate while evading the immune system. Many of these viruses, including the severe acute respiratory syndrome coronavirus 2 (SARS-CoV-2), infect the brain and may cause serious neurological symptoms through mechanisms which remain poorly understood^2–4^. Here we show that expression of either the SARS-CoV-2 spike (S) protein or p15 protein from the baboon orthoreovirus is sufficient to induce fusion between interconnected neurons, as well as between neurons and glial cells. This phenomenon is observed across species, from nematodes to mammals, including human embryonic stem cells-derived neurons and brain organoids. We show that fusion events are progressive, can occur between distant neurites, and lead to the formation of multicellular syncytia. Finally, we reveal that in addition to intracellular molecules, fusion events allow diffusion and movement of large organelles such as mitochondria between fused neurons. Our results provide important mechanistic insights into how SARS-CoV-2 and other viruses could affect the nervous system circuitries causing neurological symptoms.

## Main

Viruses from diverse families, such as rabies virus, herpes simplex virus, dengue virus, orthoreovirus, and the severe acute respiratory syndrome coronavirus 2 (SARS-CoV-2), can infect neurons ^3,5^. Viral brain infections are characterized by multiple neurological symptoms, including headache, fever, confusion, epileptic seizures, and loss of taste or smell. In more severe cases, viral brain infections can lead to encephalitis and meningitis, and potentially irreversible neuronal deficits such as paralysis and death. Most clinical symptoms originate from the death of infected neurons ^4^. However, some viruses do not kill their host cells, and the chronic neurological sequelae of these infections cannot be explained by the loss of infected neurons ^2^. Other neuropathological mechanisms must therefore underlie the progression of these viral infections leading to brain dysfunction. In non-neuronal tissues, viruses use their fusogens to fuse with host membranes and enter cells ^1^. Once expressed inside the host cell, the viral fusogens redecorate the cell membrane conferring the ability to fuse with neighboring cells. This results in the formation of multinucleated syncytia, which allow viral propagation ‘from within’, without the need for virion release into the extracellular space ^6,7^. As defined over 100 years ago by Ramon y Cajal, neurons are individual units that do not base their development or communication on cellular fusion, with the preservation of their individuality being critical for the correct function of the nervous system. It is currently unknown whether the presence of viral fusogens can cause neuronal fusion and the formation of syncytia, thereby altering the neuronal circuitry and function.

To address this question, we began by using the fusion-associated small transmembrane (FAST) fusogen p15, isolated from the baboon orthoreovirus (BRV), which infects the brain of these primates and causes meningoencephalomyelitis ^8,9^. p15 is the only viral protein required by the BRV to form a syncytium ^10^, with no receptor protein on the host cell being needed to facilitate fusion ^11,12^. To safely mimic the result of viral neuronal infection, we transfected p15 in embryonic mouse primary hippocampal neurons, and visualized the presence of fusion through the transfer of different intracellular fluorophores between neurons. Immediately after isolation, a population of neurons was co-transfected by electroporation with a plasmid containing p15 and another containing GFP; a second population of neurons was co-transfected with a plasmid containing mCherry and the empty control vector. The two neuronal populations were then plated together and maintained in culture for 7 days (7 DIV). Our results revealed that the expression of p15 was sufficient to induce neuronal fusion, as detected by the presence of neurons containing both the GFP and mCherry fluorophores (Fig. 1A first row, and B), a phenotype that was never observed when the control vector was co-transfected in the absence of p15 (Fig. 1a third row, and b). To determine whether the fluorophore diffusion was caused by the fusogenic properties of p15, we generated an inactive version of this fusogen, p15Δ21-22, in which two residues of the N-terminal of the transmembrane domain were truncated ^13^. Importantly, expression of this inactive fusogen completely abolished the fusion of neurons (Fig. 1a second row, and b).

**Fig. 1:**
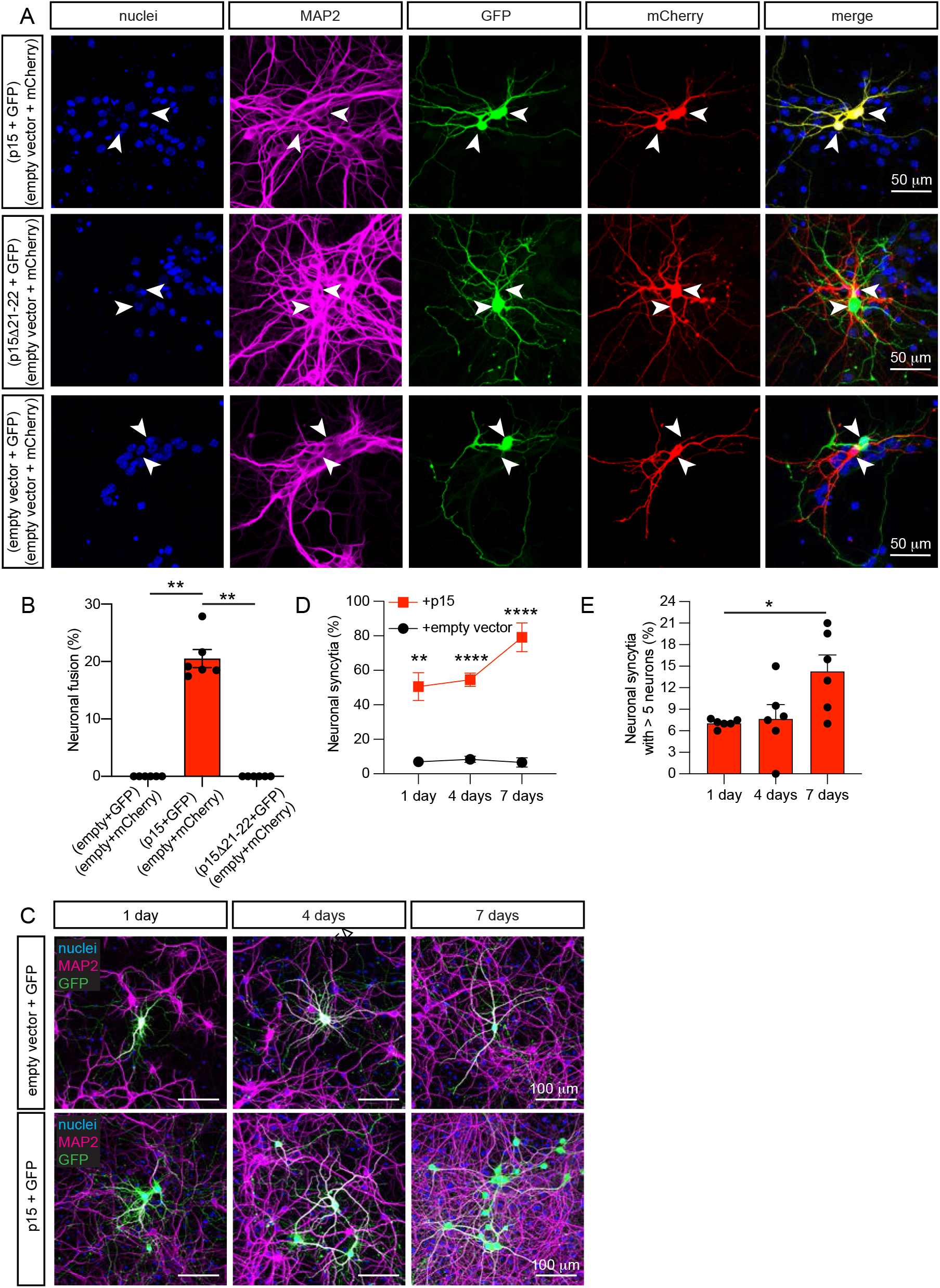
Expression of p15 induces fusion of murine neurons in culture. **a**, Representative images of fused neurons (upper row panels) identifiable with GFP (green) and mCherry (red) fluorescence appearing simultaneously in adjacent neurons (yellow in the merge panel), or non-fused control neurons (middle and lower row panels) with green and red fluorescence in adjacent neurons. Two populations of hippocampal neurons expressing either p15 and GFP, or empty vector and mCherry were cultured together for 7 days (7 DIV). In control conditions, p15 was substituted by the non-fusogenic mutant p15Δ21-22, or by the empty vector. Immunocytochemistry for nuclei (blue), neuronal MAP2 (magenta), GFP (green) and mCherry (red). **b**, Quantification of neuronal fusion as the percentage of transfected neurons that fuse (yellow) when two neurons are in proximity (≤ 200 μm). **c**, Representative images of neurons illustrating the propagation of fusion over time (upper panels). Hippocampal neurons were co-transfected at 7-10 DIV with p15 and GFP (or empty vector and GFP in control, lower panels), and were cultured for 1 day, 4 days or 7 days. Immunocytochemistry for nuclei (blue), MAP2 (magenta) and GFP (green). **d**, Quantification of neuronal syncytia as the percentage of interconnected neurons within a distance of ≤ 200 μm. **e,** Quantification of the average number of interconnected neurons per syncytium containing more than 5 neurons. Data in **b** are displayed as mean ± SEM, n > 150 neurons analyzed in 6 independent dishes from > 2 cultures, One-way ANOVA Kruskal-Wallis test followed by Dunn’s *post hoc* test in **e** comparing all groups to empty vector control. Data in **d** and **e** are displayed as mean ± SEM, n > 350 neurons analyzed in > 4 independent dishes from 4 cultures. Two-way ANOVA in **d** followed by Geisser-Greenhouse correction and the Šidák *post hoc* test comparing treatments (+ empty vector vs + p15) within each condition (days in culture). One-way ANOVA Kruskal-Wallis test followed by Dunn’s *post hoc* test in **e** comparing all groups to 1 day. \**p* <0.05, \*\**p* <0.01, \*\*\*\**p* <0.0001.

Neuronal fusion implies a temporary or permanent diffusion of cytoplasmic material between cells ^14^. To confirm that this was the case, we co-transfected neurons with p15 and the photoconvertible fluorescent protein Kaede, which shifts from green to red fluorescence upon illumination with ultraviolet (UV) light (350 – 400 nm). After identification of interconnected adjacent green fluorescent neurons, we photoconverted the green Kaede fluorophore by applying brief pulses of UV light in a small region of one neuron (donor). Newly generated red photoconverted Kaede molecules rapidly diffused to the adjacent neuron (acceptor) (Extended Data Fig. 1a and Supplementary video 1). The diffusion of red Kaede was measured as a decrease in the red fluorescence within the donor neurons (Extended Data Fig. 1b), with a concomitant increase in the red fluorescence in the acceptor neurons (Extended Data Figs. 1c, d), thereby conclusively demonstrating the existence of an active cytoplasmic bridge between p15-fused neurons. In the absence of fusion, red Kaede remained confined within the photoconverted neuron (Extended Data Fig. 1a, and 1e-g). We then asked whether the fusion bridges allowed exchange of cellular components larger than fluorescent proteins. To address this question, we used the photo-activatable mitochondrial marker mito-mPA-GFP, which allows to visualize mitochondria movement. Indeed, in fused neurons, we observed diffusion of mitochondria from donor to acceptor neurons (Extended Data Fig. 2, Supplementary Video 2).

We next asked whether the neuronal fusion induced by p15 was restricted to two adjacent neurons, or if it was a propagating event that generated syncytia with a larger number of interconnected neurons. To address this possibility, we co-transfected neurons with p15 and GFP and monitored the appearance of syncytia over a period of 7 days. The number of neuronal syncytia increased over time, appearing as clusters of interconnected GFP-positive neurons that progressively incorporated more cells (Fig. 1c-e). Moreover, through live confocal imaging performed over a period of 50 hours, we observed the progressive appearance of GFP in surrounding neurons, revealing the occurrence of fusion events (Supplementary Video 3). Fusion was observed between the somas of adjacent neurons as well as between the processes (*i.e*. dendrites and axons) of distant neurons, resulting in fusion bridges with variable lengths that could extend over hundreds of microns (Extended Data Fig. 3).

To determine if the p15-mediated neuronal fusion capacity was also conserved *in vivo*, we generated transgenic *Caenorhabditis elegans* strains in which p15 and GFP were expressed simultaneously under the control of the *mec-4* promoter (*Pmec-4::p15* and *Pmec-4::GFP*), which is active in the six mechanosensory neurons (ALM left and right, AVM, PVM and PLM left and right; Fig. 2a panel i and v). Similar to what was observed in mammalian neurons in culture, any fusion event with nearby neurons or tissues would be detectable by diffusion of GFP from the mechanosensory neurons to other cells. As predicted, we observed the appearance of additional GFP-positive cells in the head, mid body and tail of the animal, a phenomenon never observed in non-transgenic siblings or wild-type animals (Fig. 2a, b). Based on its stereotypic location and morphology, we identified ALN as the most prevalent additional GFP-positive neurons. ALN are a pair of sensory neurons located in the tail of the animal that extend their axons in close association with the axons of the ipsilateral ALM mechanosensory neurons ^15^. Other frequently GFP-positive neurons were the pair of interneurons LUA, which are located in the tail of the animal and extend anterior neurites in close proximity with those of the PLM neurons, and the mechanosensory neurons PVD, positioned in the mid-body of the animal with two extensively branched dendrites and a long axon ^15^. Despite over 90% of the GFP-positive cells being neurons, we also identified fluorescence in hypodermal cells, which form a tissue in which the PLM and ALM axons are embedded and has a glial-like function ^16,17^ (Fig. 2a, panel iv). Similar to mammalian neurons, the expression of the inactive fusogen p15Δ21-22 within mechanosensory neurons did not result in fusion with neurons or hypodermal cells (Fig. 2b). Taken together, these results indicate that the *in vivo* expression of p15 can drive the fusing of neurons with other neurons or glial cells that are located in close proximity, suggesting a possible pathomechanism of neuronal malfunction caused by orthoreovirus infection.

**Fig. 2:**
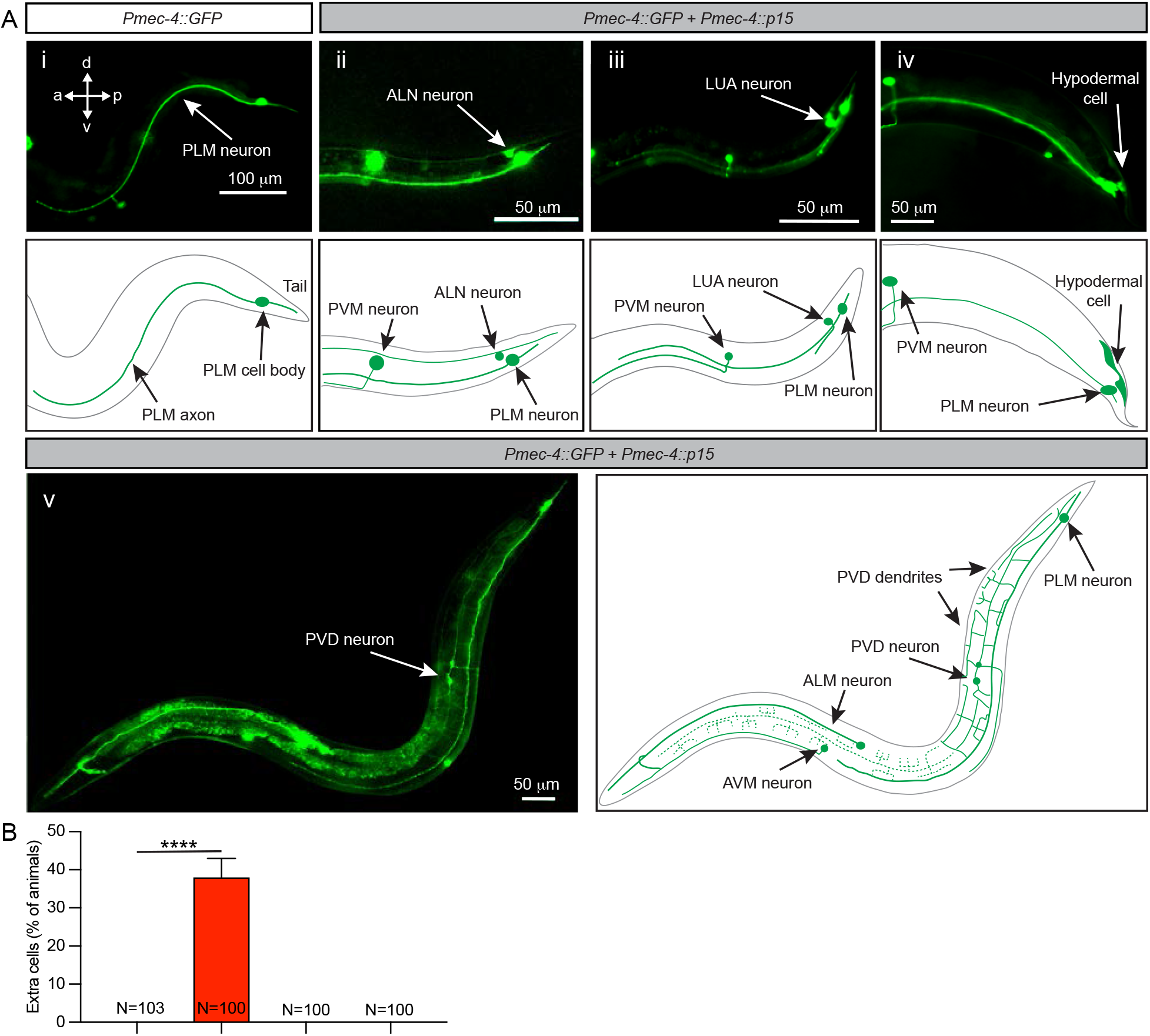
Expression of p15 induces neuronal fusion *in vivo* in *C. elegans* neurons. **a**, Representative images of animals expressing GFP in the six mechanosensory neurons (*Pmec-4::GFP*) in control conditions (i.e. no fusogen present, panel i), or when co-expressing p15 within the same neurons (*Pmec-4::GFP* + *Pmec-4::p15*) (panels ii, iii, iv, v). The anterior (a), posterior (p), dorsal (d) and ventral (v) axes are maintained through all the images and the focus is on the posterior lateral mechanosensory neuron (PLM). Representative images of the appearance of additional cells in animals expressing active p15: ALN neurons (ii), LUA neurons (iii), hypodermal (glial-like) cells (iv), and PVD neurons (v). **b**, Quantification of the percentage of animals presenting extra cells in non-transgenic siblings (-transgene), animals expressing p15, or animals expressing inactive p15 (p15Δ21-22). Data in **b** are displayed as mean ± SD, with the number of animals analyzed per condition (N) listed in the bar graph. One-way ANOVA Brown-Forsythe and Welch tests in **b** followed by Games-Howell’s *post hoc* test, comparing each mutant with its non-transgenic siblings (-transgene). \*\*\*\**p* <0.0001.

Orthoreoviruses can be transmitted from bats to humans ^18^, and SARS-CoV-2 virus is derived from bat coronaviruses ^19^. SARS-CoV-2 primarily causes a respiratory illness, but increasing evidence reveal also brain infection ^20–22^. Unlike p15, the spike S protein that functions as the fusogen of the SARS-CoV-2 virus must bind to a receptor protein, the human angiotensin-converting enzyme 2 (hACE2), located on the surface of the host cell ^23^, and uses neuropilin-1 as a co-factor to enhance infectivity ^24^. Among other tissues, hACE2 is expressed in neuronal and glial cells in the human central nervous system ^25^. Mouse neurons express mACE2 ^26^, which shares 81.86% interspecies homology with the human protein but lacks key residues for spike S binding ^23^. For this reason, to study the neuronal fusion properties of the spike S protein, both spike S and hACE2 must be expressed in murine neurons. Using a similar approach as that described for p15, we independently electroporated two neuronal populations, one with a plasmid expressing GFP plus a plasmid containing a codon-optimized spike S protein, and the other with a plasmid expressing mCherry plus a plasmid containing the hACE2 receptor. The two populations were then plated together and cultured for 7 days. The expression of the fusogen spike S and its receptor hACE2 in adjacent cells resulted in the fusion of these neurons and the mixture of the fluorophores (Fig. 3a first row, and b). Unlike p15, the presence of the specific receptor was required to initiate cellular fusion, as expression of spike S or hACE2 alone did not generate any fusion events (Fig. 3a second and third rows, and b). To determine whether the fusion of neurons was caused by the fusogenic properties of spike S, we used two fusion-inactive versions of this protein, the spike S-2P and the spike S-6P (HexaPro). First, we generated the spike S-2P variant containing two consecutive proline substitutions in the C-terminal S2 subunit ^27^. These two mutations retain the spike S in a prefusion conformation, blocking its fusion capacity ^28^. Interestingly, this mutant form has been chosen for the creation of most available vaccines against SARS-CoV-2 ^29^. The spike S-6P contains four additional proline substitutions (F817P, A892P, A899P, A942P) that further stabilize the prefusion conformation, and increase protein expression and ability to withstand heat stress ^30^. Our results indicate that none of these two versions of spike S induce neuronal fusion (Fig. 3c and d). In addition to neuron-neuron fusion, we observed neuron-glia and glia-glia fusion events if fusogen and receptor were expressed in these cell types (Extended Data Fig. 4). Neurons could fuse through their soma or via contacting neurites, forming fusion bridges of lengths that could reach over 100 microns (Extended Data Fig. 5). The functionality of these fusion bridges was evaluated using the exchange of photoconverted red Kaede fluorophores between fused neurons. Similar to what we observed with p15, photoconversion of Kaede in a small region of one fused neuron (donor) resulted in the fast diffusion of red Kaede molecules towards the other fused neuron (acceptor) (Extended Data Fig. 6). These results demonstrate that fusion through spike S and hACE2 also generates functional bridges that allow the diffusion of proteins. Importantly, the fused neurons retained their morphology, extended processes, and remained viable for the duration of the experiment.

**Fig. 3:**
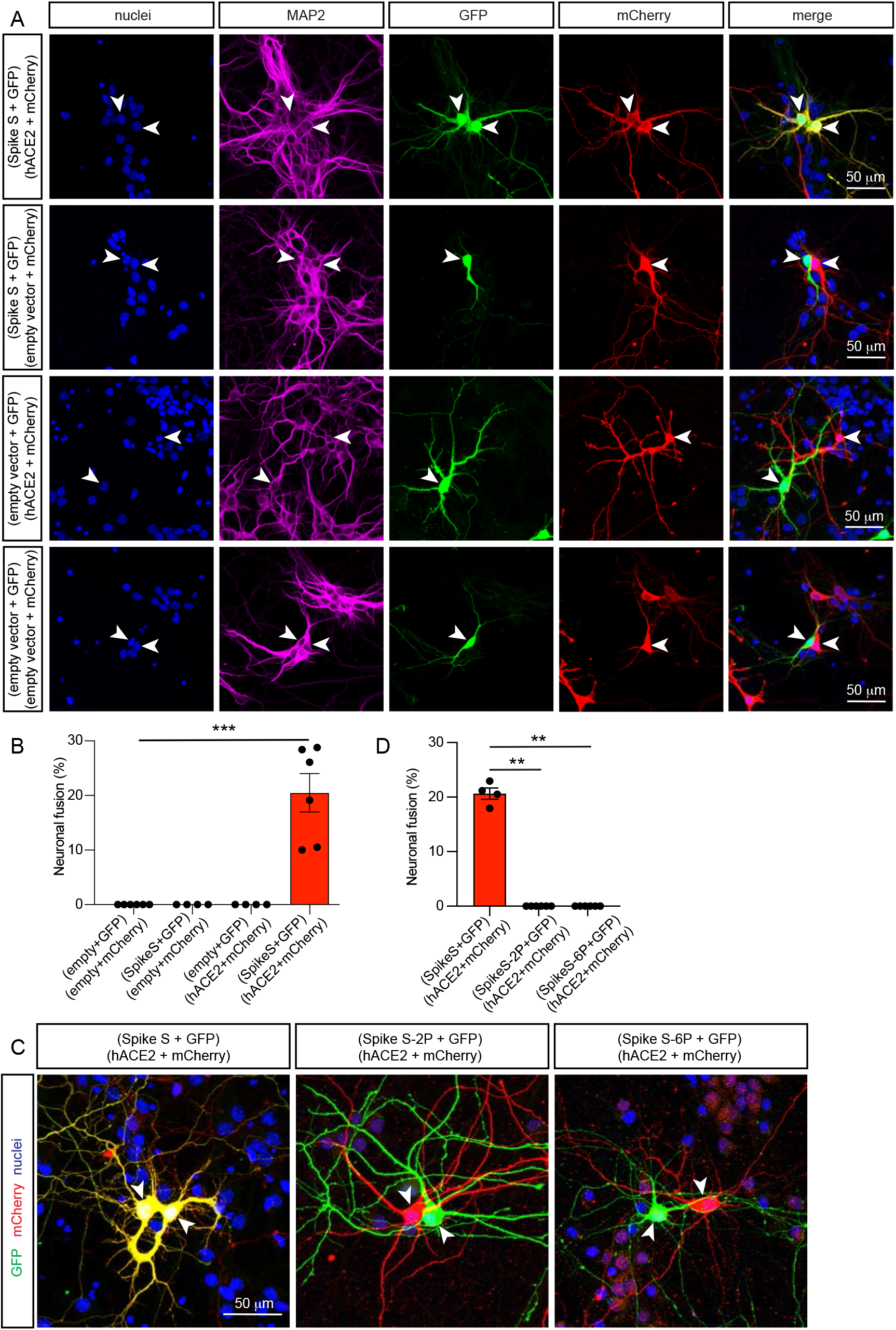
Expression of spike S and its receptor hACE2 induce fusion of murine neurons in culture. **a**, Representative images of fused neurons (first row), or non-fused control neurons (other rows). Two populations of hippocampal neurons expressing a combination of two plasmids as indicated on the left (spike S and GFP, hACE2 and mCherry, empty vector and GFP, or empty vector and mCherry) were cultured together for 7 days (7 DIV). Immunocytochemistry for nuclei (blue), MAP2 (magenta), GFP (green) and mCherry (red). Neuronal fusion only occurred (first row) when one population of neurons was transfected with spike S and GFP, and the other with hACE2 and mCherry, as visualized by the presence of GFP and mCherry in the same neurons (yellow in the merge panel). **b**, Quantification of neuronal fusion as the percentage of neurons that fuse (yellow) when two neurons are in proximity ( ≤ 200 μm). **c**, Representative images of fused neurons (first panel), or non-fused neurons (second and third panels). Two populations of hippocampal neurons expressing a combination of two plasmids as indicated above the images (spike S and GFP, hACE2 and mCherry, spike S-2P and GFP, or spike S-6P and GFP) were cultured together for 7 days (7 DIV). Immunocytochemistry for nuclei (blue), MAP2 (magenta), GFP (green) and mCherry (red). Neuronal fusion only occurred (first panel) when the full-length WT spike S protein was transfected, as visualized by the presence of GFP and mCherry in the same neurons (yellow in the panel), and not when any of the non-fusogenic mutants (spike S-2P, spike S-6P) were used (second and third panels). **d**, Quantification of neuronal fusion as the percentage of neurons that fuse (yellow) when two neurons are in proximity ( ≤ 200 μm). Data in **b** and **d** were displayed as mean ± SEM, n > 200 neurons analyzed in 4-6 independent dishes from 2 dissections, one-way ANOVA Kruskal-Wallis test in **e** followed by Dunn’s *post hoc* test comparing all groups to the group without spike S or hACE2. \*\**p* <0.01, \*\*\**p* <0.001.

Finally, we tested whether p15 and spike S have the potential to cause neuronal fusion in human embryonic stem cell (hESC)-derived neurons and brain organoids. Human neurons and organoids were transfected with a fluorophore (mCherry or GFP) with either p15, spike S or the inactive spike S-6P, and compared with their respective controls expressing fluorescence markers only. Remarkably, we observed that both p15 and full-length spike S fusogens caused extensive neuronal fusion in both human-derived systems (Fig. 4 and Extended Data Fig. 7). Importantly, the spike S-6P mutant also failed to induce cell fusion in cultured human neurons (Fig. 4b and c) and brain organoids (Fig. 4e).

**Fig. 4:**
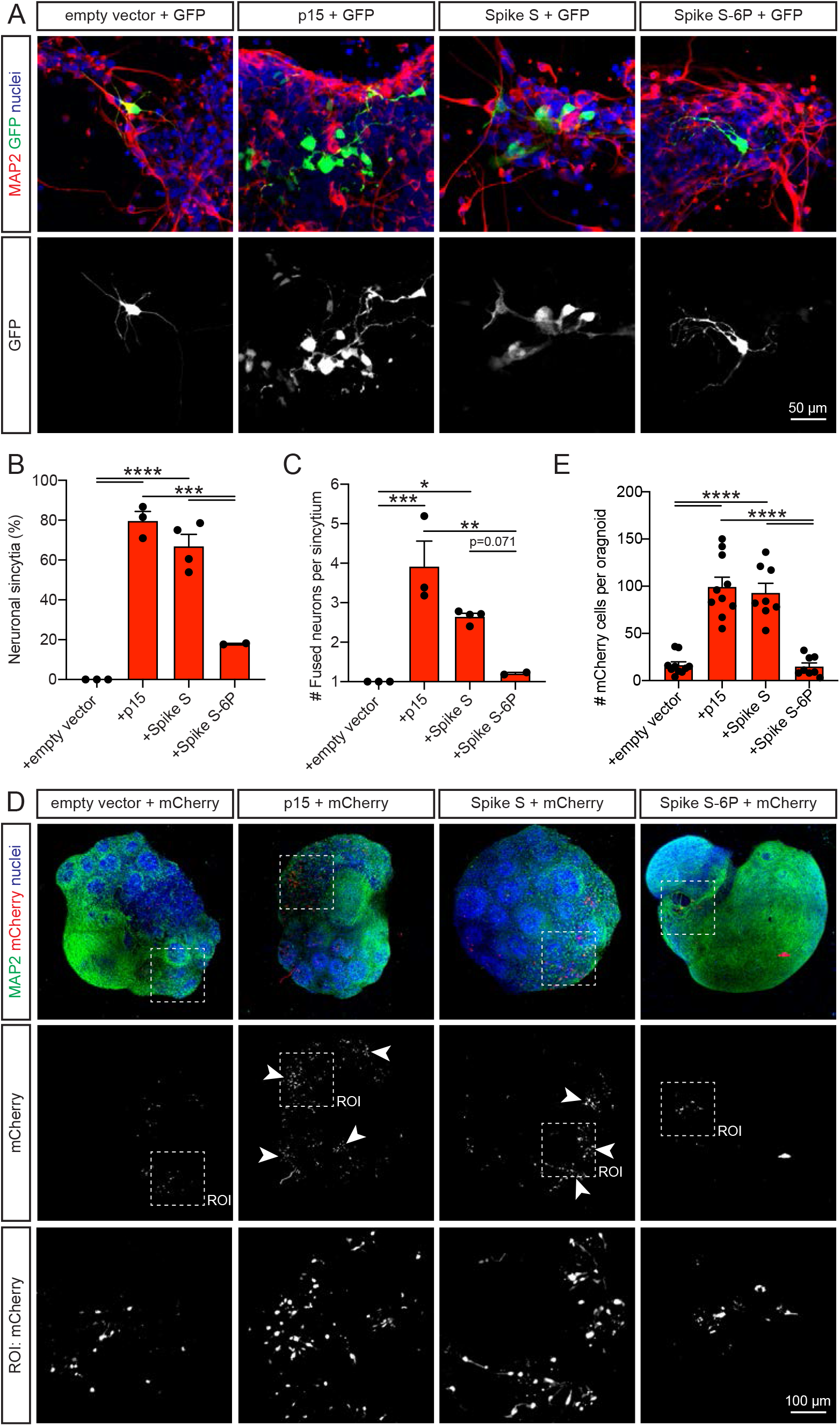
p15 and spike S induce fusion in human neurons and brain organoids. **a**, Representative images of 2D-cultured neurons illustrating fusion of cells into syncytia. Human neurons were co-transfected at 40-50 DIV with GFP and either p15, spike S or spike S-6P (or empty vector in controls), then cultured for 7 days. Immunocytochemistry for nuclei (blue), MAP2 (red) and GFP (green/white). **b**, Quantification of neuronal syncytia as the percentage of interconnected neurons within a distance of ≤ 200 μm. **c**, Quantification of the average number of interconnected neurons per syncytium containing more than one neuron. **d**, Representative images of 3D neuronal organoids illustrating fusion of cells into syncytia. Organoids were co-transfected at 43-50 DIV with mCherry and either p15, spike S or spike S-6P (or empty vector in controls), then cultured for 6 days. Immunocytochemistry for nuclei (blue), MAP2 (green) and mCherry (red/white). Regions of interest (ROIs) show higher magnification at positions indicated by broken lines. Arrowheads indicate clusters of fused neurons. **e**, Quantification of the average number of mCherry-positive cells per organoid section 6 days after transfection. Data in **b**, **c** and **e** are displayed as mean ± SEM, averages of n > 30 neurons analyzed in independent experiments for **b** and **c**, and n > 8 organoids analyzed in independent experiments for **e**. One-way ANOVA followed by Tukey *post hoc* test in **b** and **c**. \**p* <0.05, \*\**p* <0.01, \*\*\**p* <0.001, \*\*\*\**p* <0.0001.

Fused neurons can result in compromised neuronal circuitry and altered animal behavior, as previously shown for *C. elegans* chemosensory neurons that ectopically express endogenous fusogens ^31^. Our results demonstrate that neurons expressing viral fusogens acquire the capacity to fuse, potentially compromising their functional circuit properties while remaining viable. This previously uncharacterized event could explain at least some, if not most, of the neurological consequences of viral infections of the nervous system. Moreover, most of the current immunization approaches for COVID-19 are based on expressing the spike S protein in the host cells as an epitope to trigger the immune system response ^29^. These nucleic acid-based vaccines deliver the antigen encoded as mRNA, such as the Pfizer-BioNTech BNT162b2 and the Moderna mRNA-1273 vaccines ^32^, or as adenovirus-enclosed DNA, such as the Oxford-AstraZeneca ChAdOx1 nCoV-19/AZD1222 ^33^ and Johnson & Johnson Ad26.COV2.S ^28^ vaccines. The current versions of the Moderna, Pfizer-BioNTech and Johnson & Johnson vaccines encode the full-length spike S (spike S-2P) with two mutations that stabilize the pre-fusion conformation and inactivate its fusogenicity ^28,34,35^. Our findings demonstrate that it is critical to consider the fusogenic potential when designing any future vaccines in which viral fusogens are to be expressed in mammalian cells.

## Methods

### Molecular biology

Standard molecular biology methods were used. The p15 DNA sequence was obtained from the National Center for Biotechnology Information (http://www.ncbi.nlm.nih.gov/gene/). The plasmid was then designed using the software ‘A Plasmid Editor’ and the insert was generated by Integrated DNA Technologies. The *Pmec-4::p15* plasmid was constructed by subcloning p15 between Msc I and Nhe I. The *CMV::p15* plasmid was generated by subcloning *p15* into the pmaxCloning™ vector (Lonza, # VDC-1040) between Hind III and Not I. The *CMV::p15Δ21/22* plasmid was generated as previously described ^13^ by deletion of the amino acids 21 and 22 of the N-terminus of the transmembrane domain. The *CMV::SARS-CoV-2-S-2P* plasmid was generated by introducing two prolines at the 986 (K986P) and the 987 (V987P) positions of the *SARS-CoV-2-S* gene of the pCMV14-3X-Flag-SARS-CoV-2 S plasmid. Mutations were done using the QuikChangeII Site-Directed Mutagenesis Kit (*p15Δ21/22* forward primer 5’-CCACCGCCAAATGCTTTTGTTGAAAGCAGTTCTACTG-3’; *p15Δ21/22* reverse primer 5’-CAGTAGAACTGCTTTCAACAAAAGCATTTGGCGGTGG-3’; *SARS-CoV-2-S-2P* forward plasmid 5’-CCTGAGTCGCCTTGATCCGCCGGAAGCTGAAGTTC-3’; *SARS-CoV-2-S-2P* reverse plasmid 5’-GAACTTCAGCTTCCGGCGGATCAAGGCGACTCAGG-3’). Positive clones were confirmed by Sanger sequencing. The Kaede-N1 plasmid was a gift from Michael Davidson (Addgene plasmid # 54726; http://n2t.net/addgene:54726; RRID:Addgene_54726) ^36^. The mito-mPA-GFP plasmid was a gift from Richard Youle (Addgene plasmid # 23348; http://n2t.net/addgene:23348; RRID:Addgene_23348) ^37^. The pCMV14-3X-Flag-SARS-CoV-2 S plasmid was a gift from Zhaohui Qian (Addgene plasmid # 145780; http://n2t.net/addgene:145780; RRID:Addgene_145780) ^38^. The pcDNA3.1-SARS2-Spike plasmid was a gift from Fang Li (Addgene plasmid # 145032; http://n2t.net/addgene:145032; RRID:Addgene_145032) ^39^. The SARS-CoV-2 S-6P plasmid was a gift from Jason McLellan (Addgene plasmid # 154754; http://n2t.net/addgene:154754; RRID:Addgene_154754) ^30^. The pcDNA3.1-hACE2 plasmid was a gift from Fang Li (Addgene plasmid # 145033; http://n2t.net/addgene:145033; RRID:Addgene_145033) ^39^.

### Animal ethics and mouse strains

All experimental procedures using animals were conducted under the guidelines of the Australian Code of Practice for the Care and Use of Animals for Scientific purposes, and were approved by the University of Queensland Animal Ethics Committee (2019/AE000243). Wildtype (WT, C57Bl/6 background) mice were maintained on a 12 h light/dark cycle and housed in a PC2 facility with ad libitum access to food and water.

### Murine neuronal culture

Primary hippocampal neurons were obtained from mice at embryonic day E16. Isolated hippocampi were prepared as previously described ^40,41^. Briefly, 250,000 neurons were plated onto poly-L-lysine-coated 35 mm glass-bottom dishes (In Vitro Scientific) in Neurobasal medium (Gibco) supplemented with 5% fetal bovine serum (Hyclone), 2% B27, 2 mM GlutaMAX and 50 U/mL penicillin/streptomycin (Invitrogen). The medium was changed to serum-free/antibiotic-free medium 24 h post-seeding, and half the medium was changed every week.

### Human embryonic stem cell-derived neurons

Human embryonic stem cell (hESC)-derived cortical neurons were differentiated from hESCs (H9, WIC-WA09-MB-001; WiCell) using a modification of a previously published protocol ^42^. Briefly, the hESCs were maintained on Matrigel (#354230; Corning) in Essential 8 medium (#A1517001; Life Technologies). hESCs were dissociated with Accutase (#AT-104-500; Innovate Cell Technologies) at 37 °C for 1 min and seeded on AggreWell 800 (#34815; StemCell Technologies) in Essential 8 medium with ROCK inhibitor Y-27632 (10 μM, #72308, STEMCELL Technologies). After 24 h, spheroids were collected and transfered into ultra-low-attachment plates (#CLS3471; Sigma-Aldrich) with Essential 6 medium (#A1516401; Life Technologies) containing dorsomorphin (2.5 μM, #P5499, Sigma-Aldrich), SB-431542 (10 μM, #1614, Tocris) and cultured until day 13. The spheroids were then collected and placed into Matrigel (Corning)-coated 6 well-plates with DMEM/F12 (#11330057; Thermo Fisher Scientific), supplemented with 1X N2 (#17502048; Thermo Fisher Scientific). The produced neural progenitor cells (NPCs) were passaged until day 30, and then replated on poly-L-lysine (10 μg/ml #P2636; Sigma-Aldrich) and laminin (20 μg/ml #23017015; Thermo Fisher Scientific)-coated plates and maintained in Neurobasal medium (#21103049; Thermo Fisher Scientific) supplemented with B-27 (Thermo Fisher Scientific), Glutamax (#35050061; Thermo Fisher Scientific), brain-derived neurotrophic factor (BDNF, #78133; StemCell Technologies), ascorbic acid (200 μM, #72132; StemCell Technologies), GDNF (20 ng/ml, #78139.1, StemCell Technologies) and cAMP (1 μM, #A6885; Sigma-Aldrich) for at least 2 weeks.

### Brain organoids

hESC-derived brain organoids were produced as previously reported ^43^. Briefly, hESCs were incubated with Accutase (#AT-104-500; Innovate Cell Technologies) at 37 °C for 1 min. Dissociated single cells were seeded on AggreWell 800 (#34815, StemCell Technologies). 1000 single cells were added per well in Essential 8 medium supplemented with 10μM ROCK inhibitor Y-27632, and centrifuged at 100 xg for 3 min. Cultures were incubated at 37 °C and 5% CO_2_. After 24 h, the organoids were transfered into ultra-low-attachment 60ɸ dishes (#CLS3261, Sigma-Aldrich) in Essential 6 medium supplemented with 2.5 μM dorsomorphin, 10 μM SB-431542 and 2.5 μM XAV-939 (#3748, Tocris). The medium was changed every day for 5 days. On day 6, the medium was changed into Neurobasal A (#10888; Thermo Fisher Scientific) containing GlutaMax (#35050; Thermo Fisher Scientific) and B-27 without vitamin A (#12587; Thermo Fisher Scientific). The medium was supplemented with 20 ng/ml epidermal growth factor (EGF, #236-EG; R&D Systems) and 20 ng/ml basic fibroblast growth factor (bFGF, # 233-FB; R&D Systems) until day 24, when it was switched to Neurobasal A medium containing GlutaMax, B-27 without vitamin A, and supplemented with 20 ng/ml BDNF and 20 ng/ml Neurotrophin-3 (NT3, #78074; STEMCELL Technologies) until day 43. Finally, the medium was changed to Neurobasal A containing GlutaMax and B-27 without vitamin A.

### Electroporation and transfection

When required, murine neurons were electroporated using the Invitrogen Neon™ transfection system (#MPK1025, ThermoFisher Scientific) following the manufacturer’s instructions. Briefly, immediately after dissection, isolated neurons were washed twice in Ca^2+^-Mg^2+^-free PBS and resuspended in buffer R containing 1 to 2 μg of the DNA to electroporate. The conditions for the electroporation were voltage 1500 V, width 10 ms, and 3 pulses. Alternatively, the neurons were transfected after 7-12 DIV using the Lipofectamine 2000 (#11668027, Invitrogen) reagent as previously described ^40^. Following the manufacturer’s instructions, human neurons were transfected with Lipofectamine 3000 (#L3000015, Thermo Fisher Scientific) and 1.6 μg of plasmid DNA at 40 - 50 DIV; human brain organoids were transfected with Lipofectamine 3000 and 2.4 μg of plasmid DNA at 43 - 50 DIV. The transfection medium was replaced after 24 h.

### Caenorhabditis elegans strain maintenance, crosses and manipulation

Standard techniques were used for *C. elegans* maintenance, genetic crosses and manipulations ^44^. Experiments were performed on hermaphrodite animals grown at room temperature (~22 °C) on nematode growth medium plates, seeded with OP50 *Escherichia coli*. Previously generated transgenes used in this study include: *vdEx1266 [Pmec-4::p15 5ng/μl]; vdEx1268 [Pmec-4::p15 5ng/μl]*. Transgenic strains generated during this study were obtained by standard microinjection techniques ^45^, and include: *vdEx1266 [Pmec-4::p15 5 ng/μl]; vdEx1268 [Pmec-4::p15 5 ng/μl]; vdEx1417 [Pmec-4::p15Δ21/22; 15 ng/μl]; vdEx1487 [Pmec-4::p15Δ21/22; 5 ng/μl]; vdEx1489 [Pmec-4::p15Δ21/22; 5 ng/μl]*. All injection mixes had a total concentration of 100 ng/μl and contained the transgene of interest, empty pSM plasmid as a filler, and a co-injection marker for the identification of transgenic animals. As the cell-cycle transfer of extrachromosomal arrays (*vdEx*) is unstable, some animals lose the transgene. This provides an internal control for each transgenic strain, with these controls being referred to as ‘non-transgenic siblings’ or *transgene (-)*.

### Immunofluorescence staining

Murine neurons were fixed with a solution of 4% paraformaldehyde (PFA) in PBS for 10 min. Neurons were rinsed in PBS, and permeabilized with a solution of 0.1% Triton-X-100 in PBS for 10 min. They were then washed with PBS and incubated in blocking solution (5% horse serum, 1% BSA in PBS) for 1 h at room temperature. After blocking, the neurons were incubated with the primary antibodies to GFP (#AB16901, Merck Millipore), mCherry (#ab167453, Abcam) and anti-MAP2 (#188004, Synaptic Systems), diluted in primary antibody solution (1% BSA in PBS) overnight at 4 °C. They were then washed with PBS and incubated with the secondary antibodies Alexa Fluor 488 goat anti-chicken (#A-11039, ThermoFisher Scientific), Alexa Fluor 555 goat anti-rabbit (#A32732, ThermoFisher Scientific), Alexa Fluor 647 goat anti-guinea pig (#A21450, ThermoFisher Scientific) and DAPI (#62248, ThermoFisher Scientific) in the secondary antibody solution (5% horse serum in PBS) for 1 h at room temperature. Finally, the neurons were washed and mounted in Vectashield® Plus antifade mounting medium (#H-2000, Vector Laboratories). The staining protocol was slightly modified for human neurons, with fixing for 5 min, blocking with 5% (w/v) BSA (#A9418, Sigma-Aldrich) and staining for MAP2 only. Organoids were fixed with 4% PFA at room temperature or 30 min at room temperature. PBS-rinsed organoids were permeabilized with 0.5% (v/v) Triton X-100 (#X100; Sigma-Aldrich) in PBS for 30 min, followed by blocking in 5% (w/v) BSA (#A7906; Sigma-Aldrich) for 5 h. The organoids were then incubated with primary antibodies to MAP2 (#ab5392, Abcam) at 4 °C for 3 days. Following PBS washing, they were incubated with Alexa Fluor-conjugated secondary antibodies (#A11039 or #A21437, Thermo Fisher Scientific) at 4 °C for 3 days. Nuclei were counterstained with DAPI (#D1306; Invitrogen) for 30 min and washed with PBS before imaging.

### Confocal microscopy of fixed samples

The imaging for immunofluorescence microscopy was carried out on either a Zeiss Plan Apochromat 40x/1.2 NA water-immersion objective on a confocal/two-photon laser-scanning microscope (LSM 710; Carl Zeiss) built around an Axio Observer Z1 body (Carl Zeiss), equipped with two internal gallium arsenide phosphide (GaAsP) photomultiplier tubes (PMTs) and three normal PMTs for epi- (descanned) detection, or using a Zeiss Plan Apochromat 40x/1.2 NA water-immersion objective on a confocal/two-photon laser-scanning microscope (LSM 710; Carl Zeiss) and confocal microscope (LSM 880, Carl Zeiss) built around an Axio Observer Z1 body (Carl Zeiss), equipped with two internal gallium arsenide phosphide (GaAsP) photomultiplier tubes (PMTs) and three normal PMTs for epi- (descanned) detection. Both systems were controlled by Zeiss Zen Black software. Images were further processed and analyzed with FIJI-ImageJ ^46^.

### Live imaging

*C. elegans* animals were immobilized using 0.05% tetramisole hydrochloride on 3.5% agar pads. The animals were imaged with a Zeiss Axio Imager Z1 microscope equipped with a Photometrics camera (Cool Snap HQ2), and analyzed using Metamorph software (Molecular Devices) and FIJI-ImageJ. Cytoplasmic GFP was visualized with 470/80 nm excitation and 525/50 nm emission filters. Image acquisition was performed using SlideBook 6.0 (3I, Inc) and processed using FIJI-ImageJ.

For live-cell imaging of Kaede diffusion in mammalian neurons, fusogen/empty vector-Kaede-transfected neurons were bathed in imaging buffer (145 mM NaCl, 5.6 mM KCl, 2.2 mM CaCl_2_, 0.5 mM MgCl_2_, 5.6 mM D-glucose, 0.5 mM ascorbic acid, 0.1% BSA, 15 mM HEPES, pH 7.4). Neurons were visualized at 37 °C on a Zeiss Plan Apochromat 40x/1.2 NA water-immersion objective on a confocal/two-photon laser-scanning microscope (LSM 710; Carl Zeiss). For transfected neurons, a UV pulse was applied on a 5 μm x 5 μm region of interest (ROI). Photoconversion resulted in the emergence of a spot of red fluorescence that rapidly diffused through the soma and proximal dendrites of the neurons. Simultaneous green and red images were collected every 785 ms; 5 images were acquired before photoconversion and 50 images after photoconversion. Photoconversion and diffusion of the fluorophore were performed on neurons separated by 25 μm to 100 μm. In control conditions, UV light was applied first outside the neuron, 50 μm away from the soma, and then inside the soma.

Similarly, for live-cell imaging of mito-mPA-GFP diffusion on mammalian neurons, a UV pulse was applied on a 10 μm x 10 μm ROI. Photoconversion resulted in the emergence of green mitochondria that slowly moved from one neuron to the other along the fusion bridge. Green images were collected every 5 min; 72 images were acquired before photoconversion and one last image was acquired 13 h later.

### Neuron-neuron, neuron-glia and glia-glia fusion quantification in mammalian neuronal cultures

Cell-cell fusion was quantified as the percentage of pair of transfected neurons with their somas within a radius ≤ 200 μm, and that contain simultaneously GFP and mCherry. Neuron-neuron fusion is identified by two fused cells positive for MAP2 staining; neuron-glia fusion is identified by one of the two fused cells is positive for MAP2; glia-glia fusion is identified by none of the fused cells positive for MAP2. Neuronal syncytia were quantified as the percentage of interconnected neurons within a distance ≤ 200 μm.

### Statistical analysis

Results were analyzed statistically using GraphPadPrism software (GraphPad Software, Inc). The D’Agostino and Pearson test was used to test for normality. The unpaired two-tailed non-parametric Mann-Whitney U test was used for comparison of two groups, when the data were not normally distributed. For datasets comparing more than two groups, we performed one-way ANOVA Kruskal-Wallis test followed by Dunn’s *post hoc* test for multiple comparisons, one-way ANOVA Brown-Forsythe and Welch tests followed by the Games-Howell’s *post hoc* test for multiple comparisons, or two-way ANOVA followed by Geisser-Greenhouse correction and the Šidák *post hoc* test for multiple comparisons. Statistical comparisons were performed on a per-dish or a per-neuron basis. 2-4 technical replicate dishes were imaged and 2-3 independent cultures were used per condition. Each mouse dissection provided neurons from a pool of at least 6 different embryos. Values are represented as the mean ± SD or mean ± SEM. The tests used are indicated in the respective figure legends. A *p*-value below 0.05 was accepted as significant.

## Acknowledgments

We thank Rumelo Amor and his team at the Queensland Brain Institute’s Advanced Microscopy Facility for their excellent support with the microscopy, and Apurva Kumar for laboratory assistance. We also thank Perry Bartlett, Jürgen Götz and Rowan Tweedale for comments and critical appraisal on the manuscript. We thank members of the Hilliard laboratory for insightful discussions and comments. Some strains were provided by the CGC, which is funded by NIH Office of Research Infrastructure Programs (P40 OD010440), and the International *C. elegans* Gene Knockout Consortium. MAH was supported by a NHMRC Investigator Grant (APP1197860) and Project Grant (APP1068871), and an Australian Research Council (ARC) Discovery Project (160104359). MAH and FAM were supported by a NHMRC Project Grant (APP1129367) (MAH, FAM). YDK was supported by NHMRC Career Development Fellowships (1123564). LI was supported by a NHMRC Principal research Fellowship (APP1136241). The imaging was performed at the QBI Advanced Microscopy Facility, generously supported by the Australian Government through an ARC LIEF grant (LE130100078).

## Author contributions

Conceptualization: RMM, RGS, MAH

Methodology: RMM, RGS, GB, YDK, LI, MAH

Investigation: RMM, RGS, EK, ANC, MAR, ER

Visualization: RMM, RGS, EK, ANC, ER

Funding acquisition: LI, YDK, FAM, MAH

Project administration: LI, YDK, MAH

Supervision: LI, YDK, MAH

Writing – original draft: RMM, RGS, MAH

Writing – review & editing: RMM, RGS, EK, GB, FAM, YDK, LM, MAH

## Additional information

Supplementary Information is available for this paper:

Supplementary video 1

Supplementary video 2

Supplementary video 3

All data are available in the main text or the supplementary materials. Correspondence and requests for materials should be addressed to Massimo A. Hilliard.

## References

1 Martens, S. & McMahon, H. T. Mechanisms of membrane fusion: disparate players and common principles. Nat Rev Mol Cell Biol 9, 543–556, doi:10.1038/nrm2417 (2008).

2 van den Pol, A. N. Viral infection leading to brain dysfunction: more prevalent than appreciated? Neuron 64, 17–20, doi:10.1016/j.neuron.2009.09.023 (2009).

3 Song, E. et al. Neuroinvasion of SARS-CoV-2 in human and mouse brain. J Exp Med 218, doi:10.1084/jem.20202135 (2021).

4 Griffin, D. E. & Hardwick, J. M. Perspective: virus infections and the death of neurons. Trends Microbiol 7, 155–160, doi:10.1016/s0966-842x(99)01470-5 (1999).

5 John, C. C. et al. Global research priorities for infections that affect the nervous system. Nature 527, S178–186, doi:10.1038/nature16033 (2015).

6 Compton, A. A. & Schwartz, O. They might be giants: does syncytium formation sink or spread HIV infection? PLoS Pathog 13, e1006099, doi:10.1371/journal.ppat.1006099 (2017).

7 Frankel, S. S. et al. Replication of HIV-1 in dendritic cell-derived syncytia at the mucosal surface of the adenoid. Science 272, 115–117, doi:10.1126/science.272.5258.115 (1996).

8 Duncan, R., Murphy, F. A. & Mirkovic, R. R. Characterization of a novel syncytium-inducing baboon reovirus. Virology 212, 752–756, doi:10.1006/viro.1995.1536 (1995).

9 Kumar, S. et al. Reovirus-associated meningoencephalomyelitis in baboons. Vet Pathol 51, 641–650, doi:10.1177/0300985813497487 (2014).

10 Dawe, S. & Duncan, R. The S4 genome segment of baboon reovirus is bicistronic and encodes a novel fusion-associated small transmembrane protein. J Virol 76, 2131–2140, doi:10.1128/jvi.76.5.2131-2140.2002 (2002).

11 Chan, K. M. C. et al. Evolutionarily related small viral fusogens hijack distinct but modular actin nucleation pathways to drive cell-cell fusion. Proc Natl Acad Sci U S A 118, doi:10.1073/pnas.2007526118 (2021).

12 Top, D., Read, J. A., Dawe, S. J., Syvitski, R. T. & Duncan, R. Cell-cell membrane fusion induced by p15 fusion-associated small transmembrane (FAST) protein requires a novel fusion peptide motif containing a myristoylated polyproline type II helix. J Biol Chem 287, 3403–3414, doi:10.1074/jbc.M111.305268 (2012).

13 Clancy, E. K. & Duncan, R. Helix-destabilizing, beta-branched, and polar residues in the baboon reovirus p15 transmembrane domain influence the modularity of FAST proteins. J Virol 85, 4707–4719, doi:10.1128/JVI.02223-10 (2011).

14 Neumann, B., Nguyen, K. C., Hall, D. H., Ben-Yakar, A. & Hilliard, M. A. Axonal regeneration proceeds through specific axonal fusion in transected *C. elegans* neurons. Dev Dyn 240, 1365–1372, doi:10.1002/dvdy.22606 (2011).

15 Altun, Z. F., Chen, B., Wang, Z. W. & Hall, D. H. High resolution map of *Caenorhabditis elegans* gap junction proteins. Dev Dyn 244, 903, doi:10.1002/dvdy.24287 (2015).

16 Chalfie, M. et al. The neural circuit for touch sensitivity in *Caenorhabditis elegans*. J Neurosci 5, 956–964 (1985).

17 Coakley, S., Ritchie, F. K., Galbraith, K. M. & Hilliard, M. A. Epidermal control of axonal attachment via beta-spectrin and the GTPase-activating protein TBC-10 prevents axonal degeneration. Nat Commun 11, 133, doi:10.1038/s41467-019-13795-x (2020).

18 Wang, L. F. & Anderson, D. E. Viruses in bats and potential spillover to animals and humans. Curr Opin Virol 34, 79–89, doi:10.1016/j.coviro.2018.12.007 (2019).

19 Burki, T. The origin of SARS-CoV-2. Lancet Infect Dis 20, 1018–1019, doi:10.1016/S1473-3099(20)30641-1 (2020).

20 Meinhardt, J. et al. Olfactory transmucosal SARS-CoV-2 invasion as a port of central nervous system entry in individuals with COVID-19. Nat Neurosci 24, 168–175, doi:10.1038/s41593-020-00758-5 (2021).

21 Kase, Y. & Okano, H. Neurological pathogenesis of SARS-CoV-2 (COVID-19): from virological features to clinical symptoms. Inflamm Regen 41, 15, doi:10.1186/s41232-021-00165-8 (2021).

22 Zhang, B. Z. et al. SARS-CoV-2 infects human neural progenitor cells and brain organoids. Cell Res 30, 928–931, doi:10.1038/s41422-020-0390-x (2020).

23 Yan, R. et al. Structural basis for the recognition of SARS-CoV-2 by full-length human ACE2. Science 367, 1444–1448, doi:10.1126/science.abb2762 (2020).

24 Cantuti-Castelvetri, L. et al. Neuropilin-1 facilitates SARS-CoV-2 cell entry and infectivity. Science 370, 856–860, doi:10.1126/science.abd2985 (2020).

25 Khan, S. & Gomes, J. Neuropathogenesis of SARS-CoV-2 infection. Elife 9, doi:10.7554/eLife.59136 (2020).

26 Doobay, M. F. et al. Differential expression of neuronal ACE2 in transgenic mice with overexpression of the brain renin-angiotensin system. Am J Physiol Regul Integr Comp Physiol 292, R373–381, doi:10.1152/ajpregu.00292.2006 (2007).

27 Wrapp, D. et al. Cryo-EM structure of the 2019-nCoV spike in the prefusion conformation. Science 367, 1260–1263, doi:10.1126/science.abb2507 (2020).

28 Bos, R. et al. Ad26 vector-based COVID-19 vaccine encoding a prefusion-stabilized SARS-CoV-2 Spike immunogen induces potent humoral and cellular immune responses. NPJ Vaccines 5, 91, doi:10.1038/s41541-020-00243-x (2020).

29 Krammer, F. SARS-CoV-2 vaccines in development. Nature 586, 516–527, doi:10.1038/s41586-020-2798-3 (2020).

30 Hsieh, C. L. et al. Structure-based design of prefusion-stabilized SARS-CoV-2 spikes. Science 369, 1501–1505, doi:10.1126/science.abd0826 (2020).

31 Giordano-Santini, R. et al. Fusogen-mediated neuron-neuron fusion disrupts neural circuit connectivity and alters animal behavior. Proc Natl Acad Sci U S A 117, 23054–23065, doi:10.1073/pnas.1919063117 (2020).

32 Polack, F. P. et al. Safety and efficacy of the BNT162b2 mRNA COVID-19 vaccine. N Engl J Med 383, 2603–2615, doi:10.1056/NEJMoa2034577 (2020).

33 Watanabe, Y. et al. Native-like SARS-CoV-2 spike glycoprotein expressed by ChAdOx1 nCoV-19/AZD1222 vaccine. ACS Cent Sci 7, 594–602, doi:10.1021/acscentsci.1c00080 (2021).

34 Corbett, K. S. et al. SARS-CoV-2 mRNA vaccine design enabled by prototype pathogen preparedness. Nature 586, 567–571, doi:10.1038/s41586-020-2622-0 (2020).

35 Xia, X. Domains and functions of spike protein in SARS-Cov-2 in the context of vaccine design. Viruses 13, doi:10.3390/v13010109 (2021).

36 Kremers, G. J., Hazelwood, K. L., Murphy, C. S., Davidson, M. W. & Piston, D. W. Photoconversion in orange and red fluorescent proteins. Nat Methods 6, 355–358, doi:10.1038/nmeth.1319 (2009).

37 Karbowski, M. et al. Quantitation of mitochondrial dynamics by photolabeling of individual organelles shows that mitochondrial fusion is blocked during the Bax activation phase of apoptosis. J Cell Biol 164, 493–499, doi:10.1083/jcb.200309082 (2004).

38 Ou, X. et al. Characterization of spike glycoprotein of SARS-CoV-2 on virus entry and its immune cross-reactivity with SARS-CoV. Nat Commun 11, 1620, doi:10.1038/s41467-020-15562-9 (2020).

39 Shang, J. et al. Structural basis of receptor recognition by SARS-CoV-2. Nature 581, 221–224, doi:10.1038/s41586-020-2179-y (2020).

40 Joensuu, M. et al. Visualizing endocytic recycling and trafficking in live neurons by subdiffractional tracking of internalized molecules. Nat Protoc 12, 2590–2622, doi:10.1038/nprot.2017.116 (2017).

41 Fath, T., Ke, Y. D., Gunning, P., Gotz, J. & Ittner, L. M. Primary support cultures of hippocampal and substantia nigra neurons. Nat Protoc 4, 78–85, doi:10.1038/nprot.2008.199 (2009).

42 Guttikonda, S. R. et al. Fully defined human pluripotent stem cell-derived microglia and tri-culture system model C3 production in Alzheimer’s disease. Nat Neurosci 24, 343–354, doi:10.1038/s41593-020-00796-z (2021).

43 Yoon, S. J. et al. Reliability of human cortical organoid generation. Nat Methods 16, 75–78, doi:10.1038/s41592-018-0255-0 (2019).

44 Brenner, S. The genetics of *Caenorhabditis elegans*. Genetics 77, 71–94 (1974).

45 Mello, C. C., Kramer, J. M., Stinchcomb, D. & Ambros, V. Efficient gene transfer in *C. elegans*: extrachromosomal maintenance and integration of transforming sequences. EMBO J 10, 3959–3970 (1991).

46 Schneider, C. A., Rasband, W. S. & Eliceiri, K. W. NIH Image to ImageJ: 25 years of image analysis. Nat Methods 9, 671–675, doi:10.1038/nmeth.2089 (2012).

