## Supplementary Material for "The SARS-CoV-2 spike (S) and the orthoreovirus p15 cause neuronal and glial fusion"

### Extended data figures

#### Extended Data Fig. 1: Diffusion of photoconvertible fluorophores between p15-fused neurons.

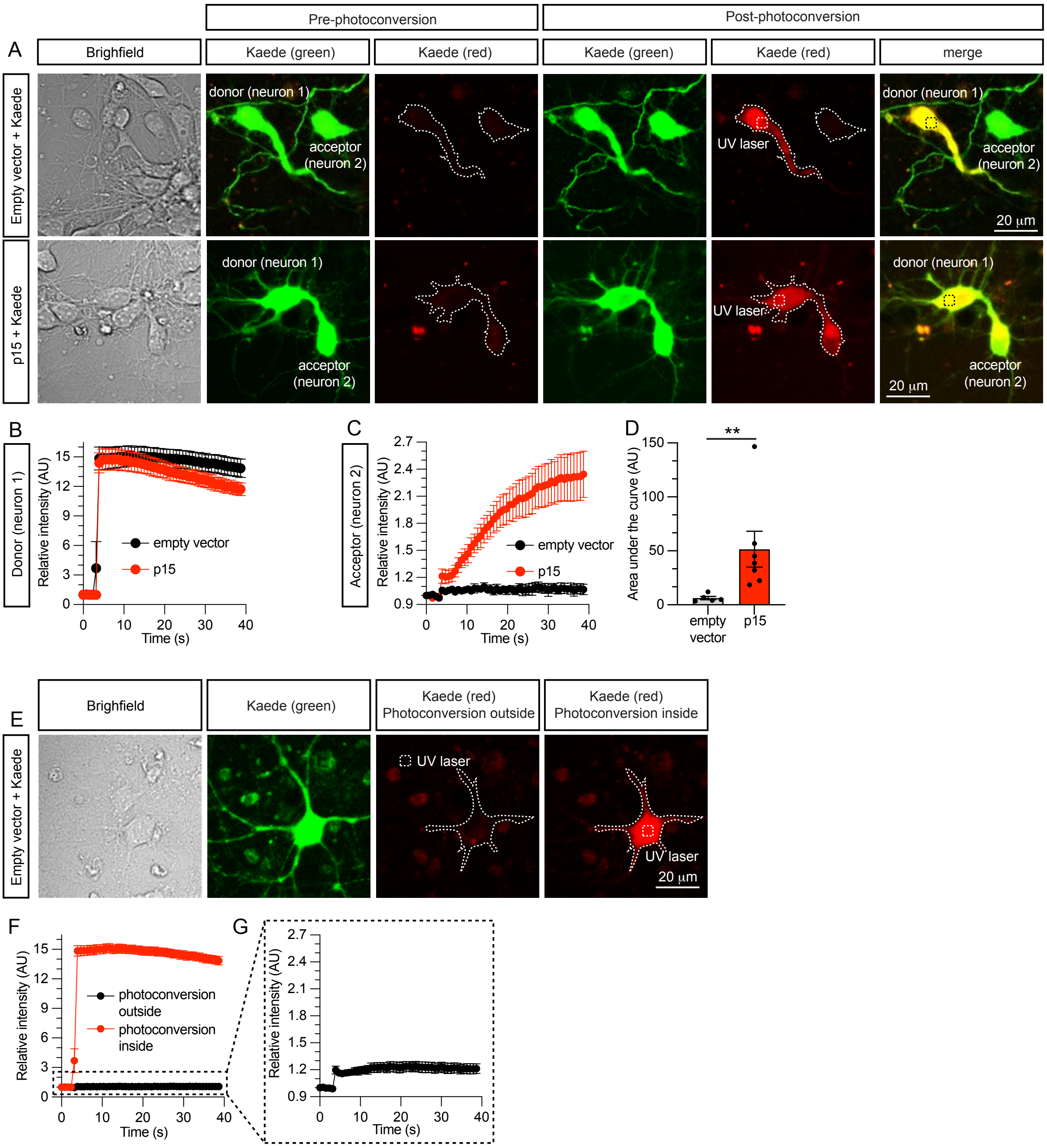


**a**, Representative images of non-fused control (empty vector) neurons (upper row panels), or fused (p15) neurons (lower row panels). Hippocampal neurons were co-transfected at DIV7 with either empty vector and Kaede (control) or p15 and Kaede. Before photoconversion (pre-photoconversion), Kaede displays a major green fluorescence emission when excited at 488 nm and negligible red emission when excited at 561 nm. After irradiation with UV light (post-photoconversion), Kaede irreversibly photoconverts to a red-emitting fluorescent protein. In the absence of neuronal fusion (upper panels), newly photoconverted red Kaede molecules cannot diffuse between adjacent cells. However, when two neurons are fused (lower panels), newly generated red photoconverted Kaede molecules rapidly diffuse from the site of photoconversion (donor-neuron 1) to the adjacent fused neuron (acceptor-neuron 2). **b**, Quantification of the decrease in the red fluorescence within the donor neurons in the absence of fusion (empty vector) or after fusion (p15). **c**, Quantification of the increase in red fluorescence within the acceptor neurons in the absence of fusion (empty vector) or after fusion (p15). **d,** Quantification of the area under the curve of the graph in **c**. **e,** Representative image of a control, non-fused neuron where the photoconversion was performed 50 μm outside (photoconversion outside), or within the same neuron (photoconversion inside). **f**, Quantification of the increase of red fluorescence after outside or inside photoconversion. **g**, Magnification of the graph obtained after outside photoconversion of control neurons. Data in **d** and **e** are displayed as mean ± SEM, n = 5 and 7 neurons from 2 independent experiments, Mann-Whitney U test. $^{**}p < 0.01$.

**Extended Data Fig. 2: Diffusion of mitochondria between p15-fused neurons.**

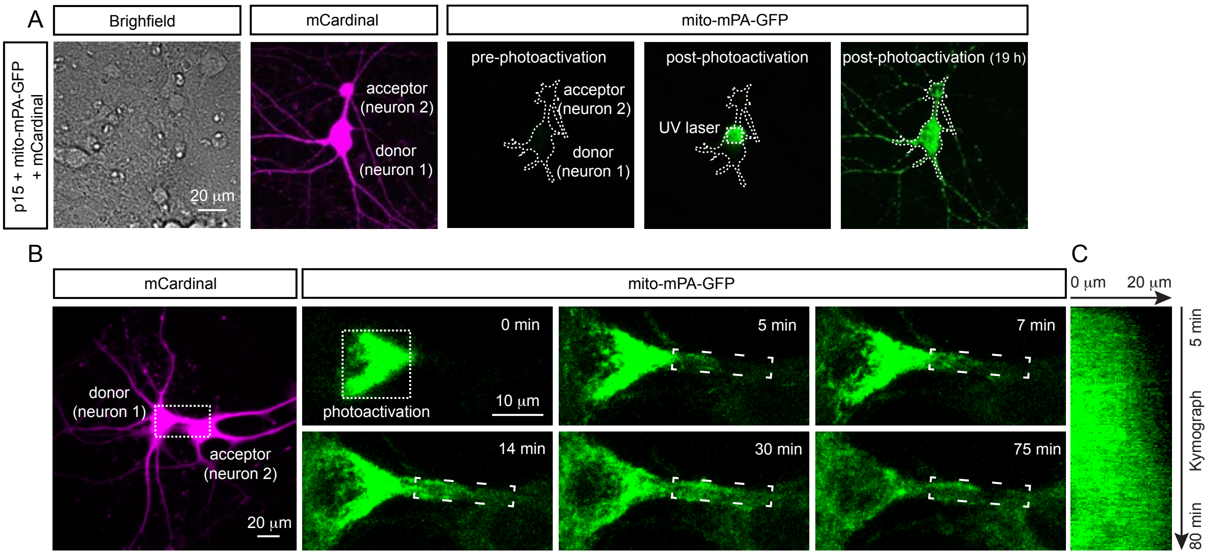


**a,** Representative images of fused hippocampal neurons co-transfected at 7 DIV with p15, the far-red fluorophore mCardinal and mito-mPA-GFP. Before photoactivation (pre-photoactivation), mito-mPA-GFP is not visible as mPA-GFP remains caged (non-fluorescent). mPA-GFP is photochemically converted to fluorescent GFP when excited with UV light. After irradiation with UV light (post-photoactivation), mPA-GFP irreversibly photoactivates to a green-emitting fluorescent protein. When two neurons are fused, newly visible mitochondria diffuse from the site of photoconversion (donor-neuron 1) to the adjacent fused neuron (acceptor-neuron 2). 19 h after photoactivation, green mitochondria are visible within the acceptor neuron. Scale bar represents 20 µm. **b,** Fused hippocampal neuron co-transfected with p15, mCardinal and mito-mPA-GFP. The boxed area is magnified in the right panels. The panels represent images of a time series showing photoactivated mito-mPA-GFP moving anterogradely along the neuronal bridge. **c,** Kymograph of mito-mPA-GFP moving along the neuronal bridge between fused neurons shown in the boxed area of the panels in **b**.

**Extended Data Fig. 3: Neuronal bridge connecting p15-fused neurons can extend over hundreds of microns.**

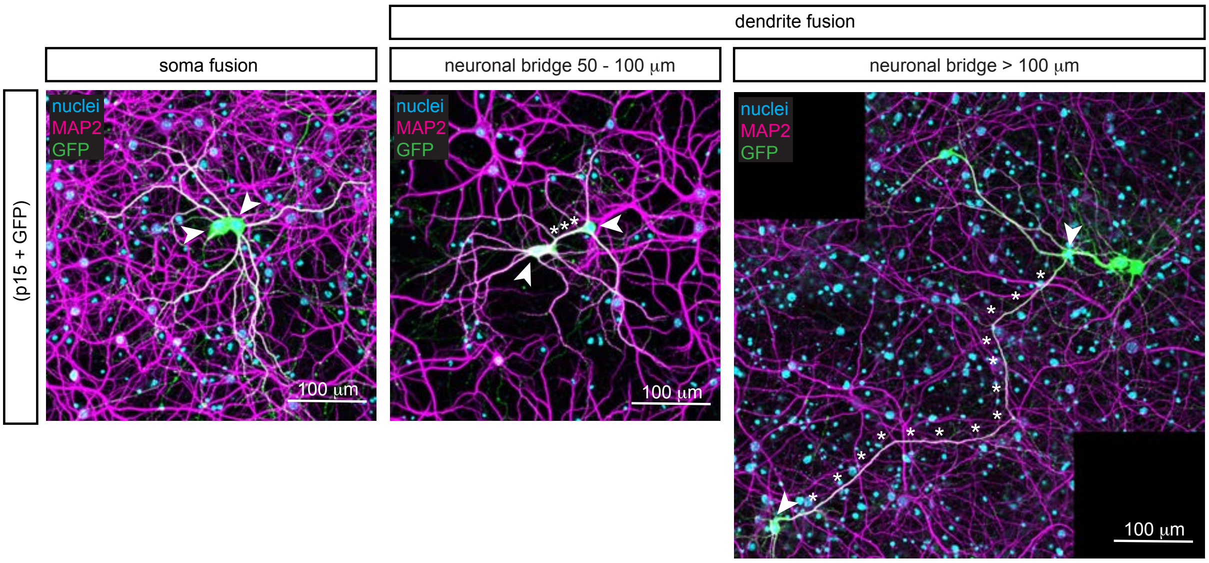


Representative images of fused neurons (arrowheads) showing fusion through the somas or through dendrites, with neuronal bridges (asterisks) having variable lengths from 50 µm to over 100 µm. Immunocytochemistry for nuclei (blue), MAP2 (magenta) and GFP (green). The third image is formed by stitching two images with overlapping regions using the FIJI-ImageJ pairwise stitching plugin.

**Extended Data Fig. 4: The expression of Spike S and its receptor hACE2 is sufficient to induce fusion of non-neuronal cells in culture.**

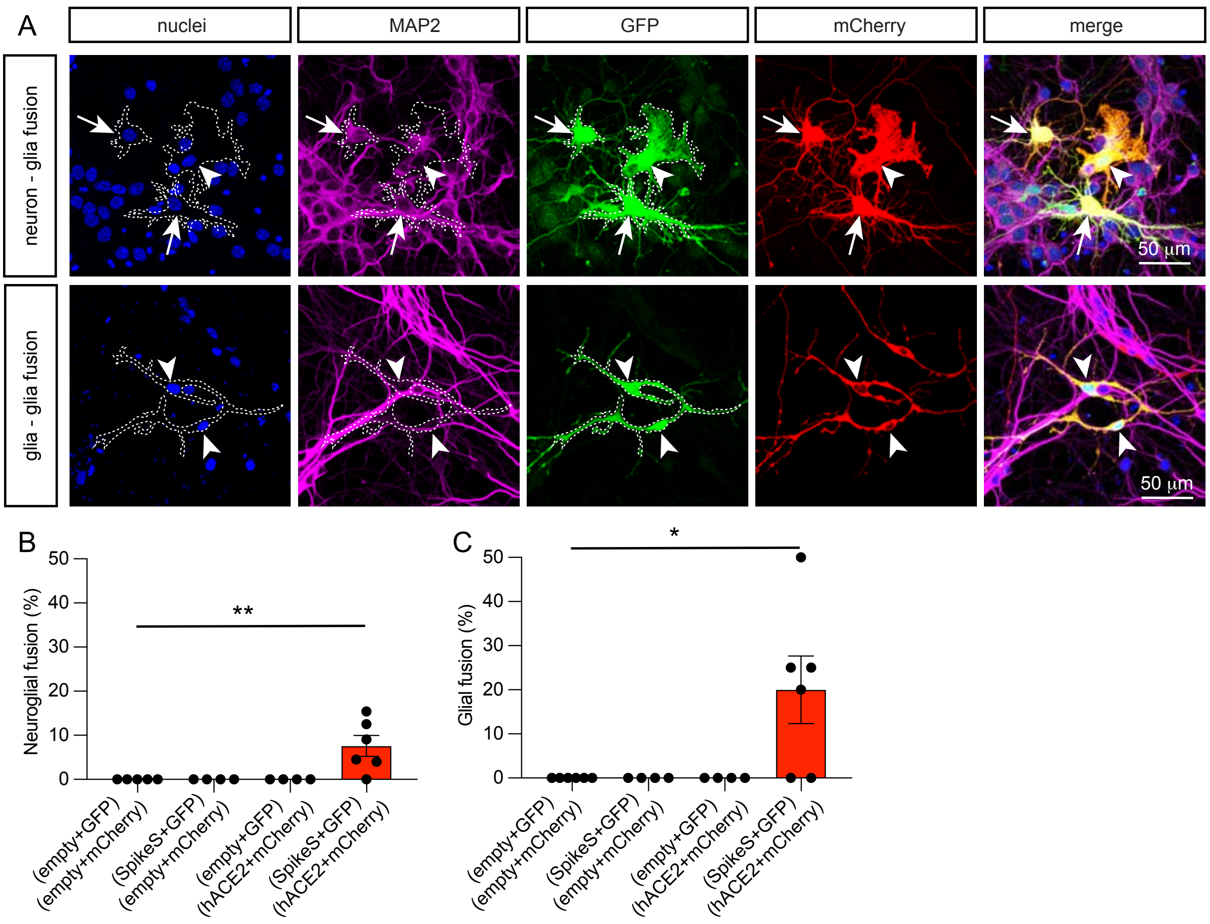


**a**, Representative images of fusion between neurons and non-neuronal glial cells (first row), or between two non-neuronal glial cells (second row). Two populations of hippocampal cells expressing a combination of two plasmids among the following, as indicated on the left: spike S and GFP, hACE2 and mCherry, empty vector and GFP, or empty vector and mCherry, and then cultured for 7 DIV. Immunocytochemistry for nuclei (blue), MAP2 (magenta), GFP (green) and mCherry (red). Neuronal fusion only occurs when one population is transfected with spike S and GFP, and the other with hACE2 and mCherry, as visualized by the presence of GFP and mCherry in the same neurons (yellow in the merge panel). Fused neurons are identified as positive for MAP2 staining (arrows in first row). Non-neuronal glial cells are identified as negative for MAP2 staining (arrowheads). **b**,

Quantification of neuron-glial fusion as the percentage of cells that fuse (yellow) when two neurons/non-neurons are in close proximity (≤ 200 μm). **c**, Quantification of non-neuronal fusion as the percentage of non-neuronal glial cells that fuse (yellow) when two non-neuronal glial cells are in direct contact. Data in **b** and **c** are displayed as mean ± SEM, n > 200 brain cells analyzed in 4-6 independent dishes from 2 dissections, one-way ANOVA Kruskal-Wallis test in **e** followed by Dunn's *post hoc* test comparing all groups to the group without Spike S nor hACE2. $*p < 0.05$, $**p < 0.01$.

**Extended Data Fig. 5: Neuronal bridge connecting SpikeS-hACE2-fused neurons can extend over hundreds of microns.**

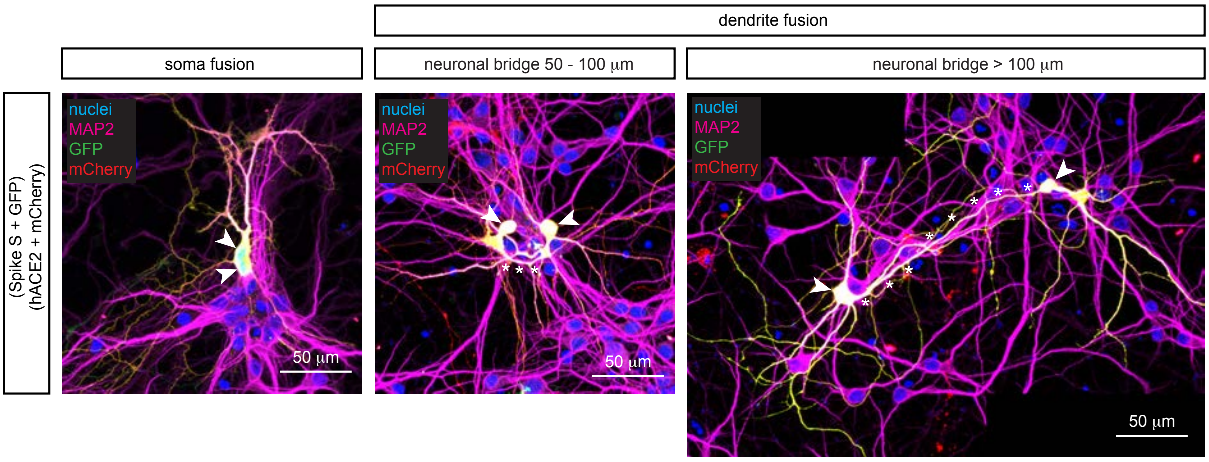


Representative images of fused neurons (arrowheads) showing fusion through the somas or through dendrites, with neuronal bridges (asterisks) having variable lengths from 50 µm to over 100 µm. Two populations of hippocampal neurons were electroporated with spike S and GFP, or hACE2 and mCherry, and then cultured together for 7 DIV. Immunocytochemistry for nuclei (blue), MAP2 (magenta), GFP (green) and mCherry (red). Neuronal fusion is identified by the presence of both GFP and mCherry in the same neurons (yellow). The third image is formed by stitching two images with overlapping regions using the FIJI-ImageJ pairwise stitching plugin.

**Extended Data Fig. 6: Diffusion of fluorophores between spike S-hACE2-fused neurons.**

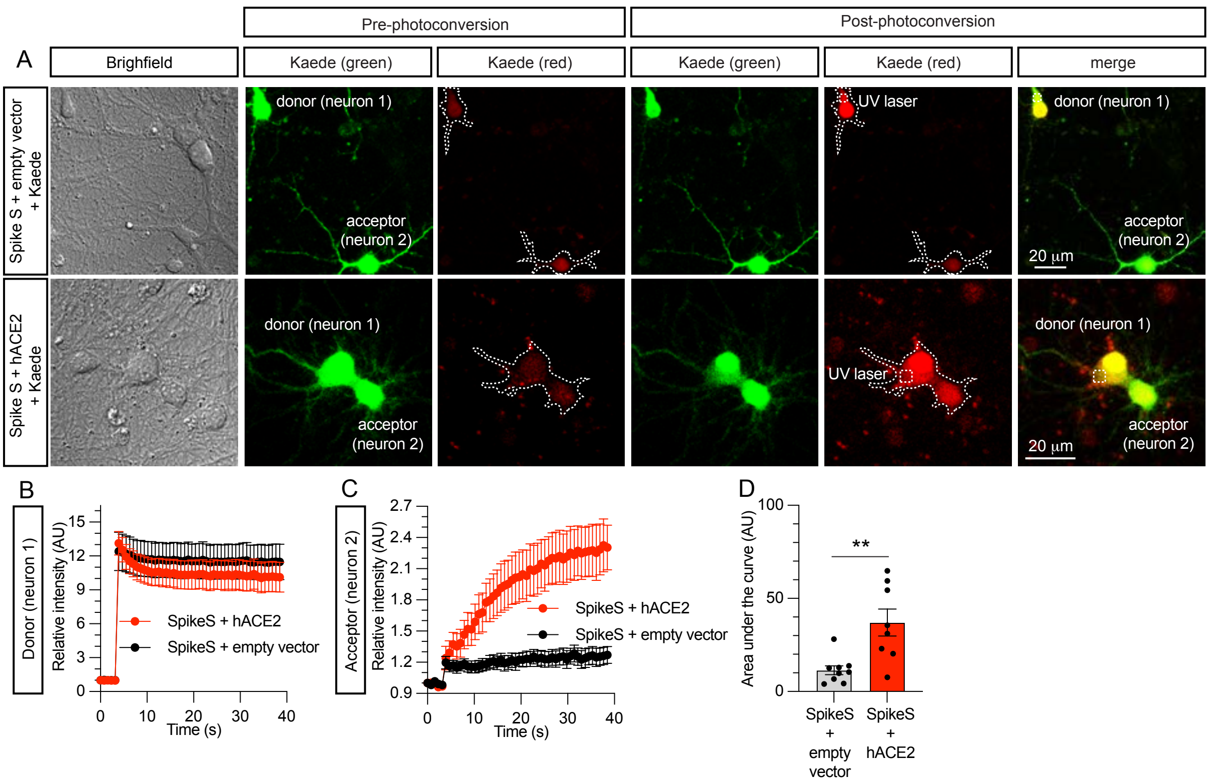


**a**, Representative images of non-fused control (spike S-empty vector) neurons (upper panels), or fused (spike S-hACE2) neurons (lower panels). Hippocampal neurons were co-transfected at 7 DIV with either spike S, empty vector and Kaede (control), or spike S, hACE2 and Kaede. Before photoconversion (pre-photoconversion), Kaede displays a major green fluorescence emission when excited at 488 nm and negligible red emission when excited at 561 nm. After irradiation with UV light (post-photoconversion), Kaede irreversibly photoconverts to a red-emitting fluorescent protein. In the absence of neuronal fusion (upper panels), newly photoconverted red Kaede molecules cannot diffuse between adjacent cells. However, when two neurons are fused (lower panels), newly generated red photoconverted Kaede molecules rapidly diffuse from the site of photoconversion (donor-neuron 1) to the adjacent fused neuron (acceptor-neuron 2). **b,** Quantification of the decrease in the red fluorescence within the donor neurons in the absence of fusion (spike S-empty vector) or after fusion (spike S-hACE2). **c,** Quantification of the increase of red fluorescence within the acceptor neurons in the absence of fusion (spike S-empty vector) or after fusion (spike S-hACE2). **d,** Quantification of the area under the curve of the graph in **c**. Data in **d** are displayed as mean ± SEM, n = 8 and 9 neurons from 2 independent experiments, Mann-Whitney U test. $**p < 0.01$.

**Extended Data Fig. 7: Temporal profiling of p15-induced cell fusion in human brain organoids.**

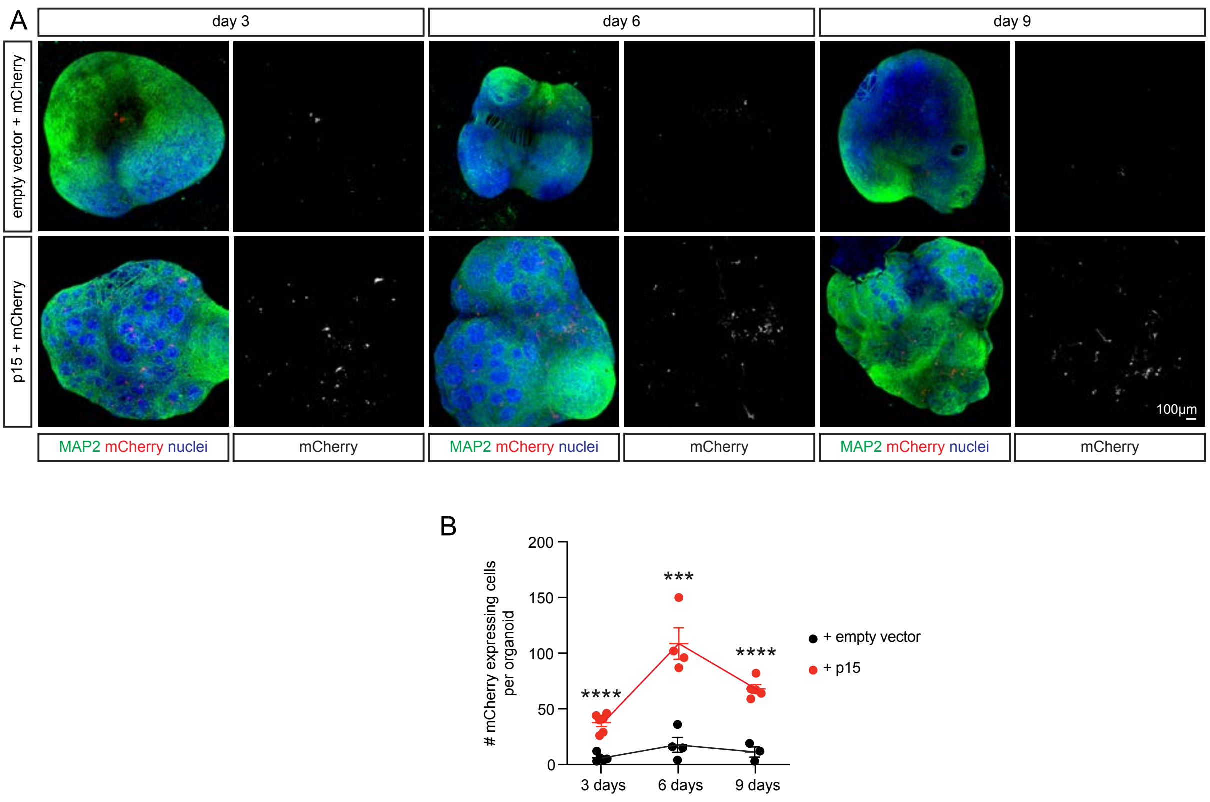


**a,** Representative images of 3D neuronal organoids illustrating fusion of cells into syncytia over time. Organoids were co-transfected at 43-50 DIV with mCherry and p15 (or empty vector in controls) and were then cultured for 3, 6 or 9 days. Immunocytochemistry for nuclei (blue), MAP2 (green) and mCherry (red/white). **b,** Quantification of the average number of mCherry-positive cells per organoid 3, 6 and 9 days after transfection. Data in **b** are displayed as mean ± SEM. One-way ANOVA followed by Tukey *post hoc* test in **b**. ***$p$ <0.001, ****$p$ <0.0001.

#### Supplementary Information

**Supplementary video 1: Diffusion of photoconverted red Kaede molecules between p15-fused neurons.** Movie at 20 frames per second (fps), with frames being acquired every 785 ms. 5 frames were acquired before photoconversion and 50 frames after photoconversion. Photoconversion was performed by applying a UV pulse on a 5 μm x 5 μm ROI (white square). Scale bar represents 20 μm.

**Supplementary video 2: Diffusion of mitochondria between p15-fused neurons.** Movie at 5 fps, with frames being acquired every 5 min. Images were recorded 10 min after mito-mPA-GFP photoactivation.

**Supplementary video 3: Appearance of newly p15-fused neurons.** Movie at 10 fps, with frames being acquired every 30 min. Images were recorded 24 h after transfecting hippocampal neurons with p15 and GFP. Each frame represents the maximum intensity projection, with GFP intensity adjusted to facilitate the visualization of newly appearing GFP-positive neurons. Scale bar represents 50 μm.
